# 3′-UTR shortening contributes to subtype-specific cancer growth by breaking stable ceRNA crosstalk of housekeeping genes

**DOI:** 10.1101/601526

**Authors:** Zhenjiang Fan, Soyeon Kim, Yulong Bai, Brenda Diergaarde, Steffi Oesterreich, Hyun Jung Park

## Abstract

Shortening of 3′UTRs (3′US) through alternative polyadenylation (APA) is a post-transcriptional mechanism that regulate expression of hundreds of genes in human cancers. In breast cancer, different subtypes of tumor samples, such as estrogen receptor positive and negative (ER+ and ER-), are characterized by distinct molecular mechanisms, suggesting possible differences in the post-transcriptional regulation between the subtype tumors. In this study, based on the profound tumorigenic role of 3′US interacting with competing-endogenous RNA (ceRNA) network (3′US-ceRNA effect), we hypothesize that the 3′US-ceRNA effect drives subtype-specific tumor growth. However, we found that the subtypes are available in different sample size, biasing the ceRNA network size and disabling the fair comparison of the 3′US-ceRNA effect. Using normalized Laplacian Matrix Eigenvalue Distribution, we addressed this bias and built the tumor ceRNA networks comparable between the subtypes. Based on the comparison, we identified a novel role of housekeeping (HK) genes as stable and strong miRNA sponges (sponge HK genes) that synchronize the ceRNA networks of normal samples (adjacent to ER+ and ER- tumor samples). We further found that distinct 3′US events in the ER- tumor break the stable sponge effect of HK genes in a subtype-specific fashion, especially in association with the aggressive and metastatic phenotypes. Knockdown of NUDT21, a master 3′-UTR regulator in HeLa cells, confirmed the causal role of 3′US-ceRNA effect repressing HK genes for tumor growth. In this study, we identified 3′US-ceRNA effect on the sponge HK genes for subtype-specific growth of ER- tumors.

## INTRODUCTION

Approximately, 70% of human genes contain multiple polyadenylation (polyA) sites in the 3′-untranslated region (3′-UTR) [1]. Through alternative polyadenylation (APA) during transcription, messenger RNAs (mRNA) from the same gene can have various 3′-UTR lengths. Since the 3′-UTR contains regulatory regions including microRNA (miRNA) target sites, mRNAs with shortened or lengthened 3′-UTRs may diversify the regulation landscape, for example miRNA binding landscape. In human cancer, 3′-UTR lengthening (3′UL) has been associated with cell senescence [2] with implications for tumor-associated processes, such as cell cycle inhibition, DNA damage/repair process, and tumor suppression [3]–[6]. Widespread 3′-UTR shortening (3′US) has been reported for diverse types of human cancer [1]. Further, 3′US events add prognostic power beyond common clinical and molecular covariates in cancer patients [7] and are associated with drug sensitivity in cancer cell lines [8]. These results suggest that APA events, both 3′-UTR shortening and lengthening, play important roles in cancer etiology and treatments.

The 3′-UTR is also implicated in competing-endogenous RNA crosstalk (ceRNA) [9]. CeRNAs co-regulate each other RNAs through competing for binding miRNAs. In diverse types of cancer, ceRNA regulation involves established oncogenes and tumor suppressor genes [10] and facilitates molecular pathway interactions for tumorigenesis [11]. When 3′-UTR shortening genes lose miRNA target sites on their 3′-UTRs and do not sequester the miRNAs, the associated miRNAs bind to the 3′-UTR of the ceRNA partners. As a result, 3′-UTR shortening disrupts ceRNA crosstalk (3′US-ceRNA effect) for growth in diverse types of cancer, including breast cancer [12]. In a recent study, we showed that this 3′US-ceRNA effect promotes tumor growth independent of potential confounding factors, such as somatic mutation status (SNPs and small INDELs), tumor purity, immune cell infiltration, cell proliferation, or miRNA biogenesis and expression [13].

Breast cancer can be classified into two major subtypes based on the presence or absence of estrogen receptor (ER) [14]. Estrogen receptor positive (ER+) breast tumors grow in the presence of the hormone estrogen. So, ER+ cancers can be treated with endocrine therapy which blocks ER activity or depletes estrogen levels. On the other hand, estrogen receptor negative (ER-) breast tumors have unique growth mechanism due to absence of the estrogen receptor. The unique growth mechanism of ER- tumors makes it difficult to treat ER- breast cancer that has a worse prognosis than ER+ [15] with a more aggressive phenotype [16], [17]. Based on the profound tumorigenic effect of 3′US-ceRNA [12], we hypothesize that 3′US-ceRNA effects specific to ER- breast tumors contribute to the unique growth mechanism. In this study, we tested this hypothesis by addressing a quantitative challenge due to different sample sizes between ER+ and ER- breast tumor samples. As a result, we identified a novel subset of housekeeping (HK) genes (sponge HK) effectively sponging miRNAs to synchronize the ceRNA networks in normal samples (adjacent to the subtype tumor samples). Further, we showed that the 3′US-ceRNA effects repress the sponge HK genes, leading to subtype-specific tumor growth. In ER- breast tumor, this subtype-specific tumor growth is associated with aggressive and metastatic phenotypes of ER- tumors, attributing its unique grow mechanism partially to subtype-specific 3′US-ceRNA effects.

## MATERIALS AND METHODS

### TCGA breast tumor RNA-seq data and identification of breast cancer subtypes

Quantified gene expression files (RNASeqV1) for primary breast tumors (TCGA sample code and their matching solid normal samples (TCGA sample code 11) were downloaded from the TCGA Data Portal[18]. We used 97 breast tumor samples that have matched normal tissues, which were further categorized into 77 estrogen receptor positive (ER+) and 20 estrogen receptor negative (ER-). For ER+ and ER-, we collected both normal (ER+ normal and ER- normal) and tumor (ER+ tumor and ER- tumor) samples. A total of 10,868 expressed RefSeq genes (fragments per kilobase of transcript per million mapped reads (FPM) ≥ 1 in > 80% of all samples) were selected for downstream analyses.

### Selection of miRNA target sites

Predicted miRNA-target sites were obtained from TargetScanHuman version 6.2[19]. Only those with a preferentially conserved targeting score (Pct) more than 0 were used[7]. Experimentally validated miRNA-target sites were obtained from TarBase version 5.0[20], miRecords version 4[21] and miRTarBase version 4.5[22]. The target sites found in indirect studies such as microarray experiments and high-throughput proteomics measurements were filtered out [23]. Another source is the microRNA target atlas composed of public AGO-CLIP data[24] with significant target sites (q-value < 0.05). The predicted and validated target site information was then combined to use in this study.

### Statistical significance of Pearson correlation coefficient

The implementation of the Pearson r function is provided by a python package, SciPy, and available at https://scipy.org/, which returns the calculated correlation coefficient and a 2-tailed p-value for testing non-correlation. The Pearson correlation coefficient measures the linear relationship between two variables (e.g. gene X and gene Y) and when the two covariates follow binormal distribution, we can assume that their Pearson’s correlation follows student t distribution. The p-value is calculated by three steps: 1) calculating the value of the Pearson’s correlation *t*, 2) defining the degree of freedom *df* (*N-2*, where *N* is the sample size), 3) getting the probability of having *t* or more extreme than *t* from a Student’s t-distribution with the degrees of freedom *df*. We used hypergeometric test in Scipy to estimate significant of miRNA binding site overlap between genes.

### Housekeeping, transcription factor and tumor-associated genes

Housekeeping genes are required for the maintenance of basic cellular functions that are essential for the existence of a cell, regardless of its specific role in the tissue or organism. Generally, housekeeping (HK) genes are expected to be expressed at relatively constant rates in most non-pathological situations[25]. We used 3,804 HK genes defined in RNA-Seq data for 16 normal human tissue types: adrenal, adipose, brain, breast, colon, heart, kidney, liver, lung, lymph, ovary, prostate, skeletal muscle, testes, thyroid, and white blood cells[26].

Transcription factors (TFs) play an important role in the gene regulatory network. We used 2,020 TF genes defined in TFcheckpoint database[27], in which TF information is collected from 9 different resources followed by manual inspections for sequence-specific DNA-binding RNA polymerase II TF.

The tumor-suppressor genes and oncogenes were defined by the TUSON algorithm from genome sequencing of > 8,200 tumor/normal pairs[28], in particular residue-specific activating mutations for oncogenes and discrete inactivating mutations for tumor-suppressor genes. TUSON computationally analyzes patterns of mutation in tumors and predicts the likelihood that any individual gene functions as a tumor-suppressor gene or oncogene. We used 466 oncogenes and 466 tumor suppressor genes at the top 500 in each prediction (after subtracting 34 genes in common).

### Building subtype ceRNA networks

For each of the breast cancer data (ER+ normal, ER+ tumor, ER- normal, and ER- tumor) that we defined above, we constructed a ceRNA network based on microRNA (miRNA) target site share and expression correlation[12], [29]. The same miRNA target site information was determined regardless of the subtypes, resulting into a miRNA target site share network (based on FDR > 0.05 in hypergeometric test with miRNA target site information). And given the same miRNA target site share network, the expression correlation information for each subtype will select ceRNA network edges for each subtype.

We first constructed the ER+ normal reference ceRNA network by applying a traditional correlation cutoff (>=0.6) on the miRNA target site share network. Then, to identify ER- normal ceRNA network comparable to ER+ normal reference ceRNA network, we applied different correlation cutoff values (0 to 1 with a step size of 0.01) on the miRNA target site share network for ER- normal samples, and select the correlation cutoff values that makes ER- normal ceRNA network most similar to ER+ normal reference ceRNA network. To estimate topological similarity, we employed normalized Laplacian Matrix Eigenvalue Distribution that discovers ensembles of Erdős–Rényi graphs better than other metrics such as Sequential Adjacency or Laplacian[30]. After identifying the ER+ normal reference network and the corresponding ER- normal network, we used the same cutoffs (0.6 for ER+ subtypes and 0.68 for ER- subtypes) to construct the ER+ tumor network and the ER- tumor network, respectively.

### Estimating topological similarity

To identify the structural equivalence between two networks, we employed spectral analysis not only to identify the structural similarities, but also to track down the underlying dynamic behavior changes between them. Spectral clustering on networks uses the eigenvalues of several matrices, such as adjacency matrix, the Laplacian matrix, the normalized Laplacian matrix. In this research, we used the normalized Laplacian matrix since it involves both the degree matrix and adjacency matrix, where the degree matrix can identify the node related equivalence of networks and the adjacency matrix can capture the structural equivalence of networks. Another very important reason of using the normalized Laplacian eigenvalue matrix is that it is more sensitive to small changes because it considers more information^17^.

For network G, the normalized Laplacian of G is the matrix:

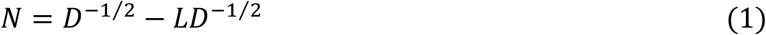

where L is the Laplacian matrix of G and D is the degree matrix. The Laplacian matrix L is defined as: *L* = *D* − *A*, where A is the adjacency matrix of G.

In N, each of its entry elements is given by:

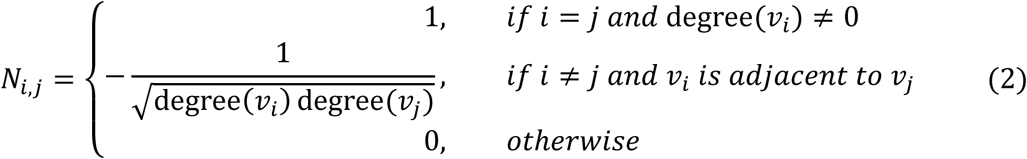

where degree (vertex v) is the function that return the degree of the vertex v.

To assess how close two eigenvalue distributions of network G_1_ and G_2_ are, we used the Kolmogorov–Smirnov test (KS test), which is defined as:

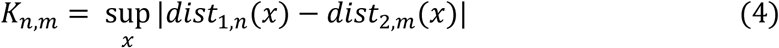

where *dist*_1,*n*_*and dist*_2,*m*_ are the empirical distribution functions of the first and the second eigenvalue distribution respectively, and 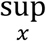 is the supremum of the set of distances.

By using the normalized Laplacian Matrix and KS test, ER+ normal reference network 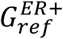 is compared with a ER- normal subnetwork with a particular correlation cutoff *i 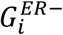* in the following three steps:

1. Compute the normalized Laplacian metrics 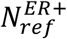 and 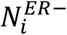 from 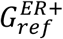 and 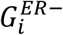 respectively.
2. Compute the eigenvalues 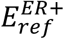 and 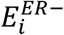 from 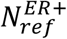 and 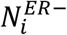 respectively.
3. Compute the KS statistic between 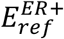 and 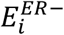.

The third step test the null hypothesis that eigenvalues 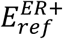 and 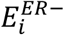 are drawn from the same continuous distribution. If the two-tailed p-value returned by the KS test is high, then we cannot reject the hypothesis that 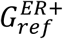 and 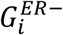 are the same network. In another word, the higher the p-value is, the more similar 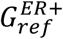 and 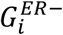.

## RESULTS

### Widespread 3′-UTR shortening and lengthening events for ER+ and ER-

To identify subtype-specific APA genes, we first identified 77 ER-positive (ER+) and 20 ER- negative (ER-) sample pairs (breast tumor and the adjacent normal samples) from 97 sample pairs available in TCGA (see Methods). Then, we identified <u>3′U</u>TR <u>s</u>hortened (3′US) and <u>3′U</u>TR lengthened (3′UL) genes (tumor vs. normal) using DaPars [7] in each subtype. We found that the ER+ and ER- sample pairs have similar numbers of total 3′US genes and 3′UL genes (**Fig. 1A**). However, 3′US genes are more recurrent (occurring in > 20% of the tumor samples [7]) in both the subtype tumors (**Fig. 1B, C** e.g. P=5.0×10^−5^ for ER+). Further analyses showed that 3′US and 3′UL play distinct roles in the subtypes. First, the recurrent 3′US and 3′UL genes show little overlap (1 and 13 genes in common, P=1.27e^−6^ and P=3.97e^−9^, respectively, **Fig. 1B, C**). Second, the number of 3′UL events is not correlated with that of 3′US events across the tumor samples (P=0.35 for ER+ and P=0.61 for ER-, **Fig. 1D, E**). Third, Ingenuity Pathway Analysis (IPA) (**S. Table 1, S. Fig. 1**) shows that the recurrent 3′US and 3′UL genes are enriched for distinct sets of molecular pathways. The IPA analysis further suggests that 3′UL or 3′US genes themselves have limited roles for cancer overall, since a small number of pathways are significantly (P<10^−2^) enriched for them and at most a couple of them are “cancer” pathways (one for 3′UL in ER+ and two for 3′US in ER- with keyword “cancer”). Based on the profound tumorigenic role of 3′US in its interaction with ceRNAs (3′US-ceRNA effect) [12], we hypothesize that 3′US-ceRNA effect, not 3′US cis effect, promotes ER- specific tumor growth.

**Figure 1.**
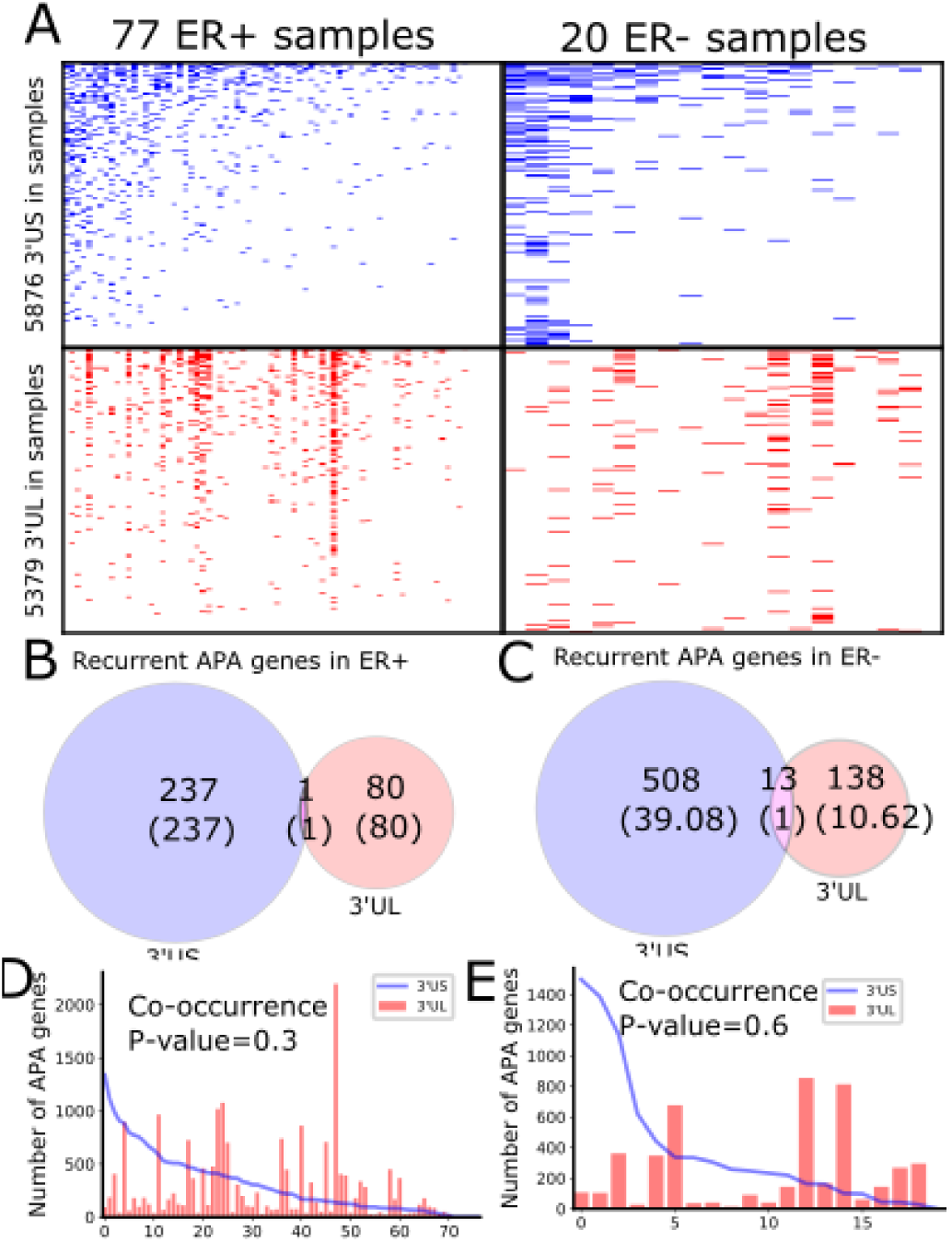
Global APA events distinct for ER+ and ER-. (A). Heatmaps showing the genes with 3′US (top panel) or 3′UL (bottom panel) in ER+ samples (left column) or ER- samples (right column). Overlap of the recurring (>20% in tumor samples) 3′US and 3′UL genes in ER+ (B) and ER- (C). In ER+ (D) and ER- (E), the number of APA genes (3′US in line and 3′UL in red bar) in the tumor-normal sample pairs ordered as in Fig. A.

### Two-step Pairwise Normalization of ER+ and ER- ceRNA network

We previously identified the 3′US-ceRNA effect in the ceRNA network [12]. To identify the 3′US-ceRNA effect specific to ER- tumors, we aim to build ceRNA networks for ER- and ER+ tumors and compare them. Computationally, ceRNA gene pairs in the networks are those that share a significant number of miRNA target sites and are co-expressed [12], [29]. However, using the common co-expression cutoff (e.g., Pearson’s ρ > 0.6) will inflate the number of edges for ER- (160,687 in ER- normal vs. 88,275 in ER+ normal, **S. Fig. 2A**). To test if this inflation is attributable to the sample size difference, we built the ceRNA network 100 times from different numbers of (20, 40, 60, and 75) normal subsamples from ER+ tumors based on the same co-expression cutoff (**S. Fig. 2B)**. In general, the number of edges in the ceRNA networks increases as the subsample size decreases. Especially, when the same number of subsamples (20) is used, the number of edges in the subsampled networks becomes closer to the inflated number.

Since the network size difference is due to sample size, one might want to subsample ER+ normal samples to match the number of samples for ER- (n=20). To assess this solution, we subsampled 20 ER+ normal samples 100 times, built a ceRNA network for each subsample, and collected all the edges (916,999) across the networks. Then, we checked how many times each edge occurs across the 100 subsampled networks. We found that the subsampled ceRNA networks do not keep topological consistency within them, as 22.1% (202,997) of the edges are shared by less than the 20 ceRNA networks (**S. Fig. 2C**). Then, one might want to build the ER- ceRNA network using the co-expression cutoff with the same statistical significance to ER+ (0.91, P∼=10^−8.2^, **S. Fig. 2D**). However, our previous simulation (**S. Fig. 2A)** showed that it will drastically deflate the number of edges. We addressed this issue in the following way. First, we built the reference network from normal samples of larger size (ER+) using the common correlation cutoff (Pearson’s ρ > 0.6). Since normal samples should have similar molecular dynamics between ER+ and ER-, we sought to find the co-expression cutoff for ER- normal network that yields the most topological similarity to the ER+ reference network. To estimate topological similarity, we employed a normalized Laplacian Matrix Eigenvalue Distribution that discovers ensembles of Erdős–Rényi graphs better than other metrics, such as Sequential Adjacency or Laplacian [30] (see Methods). While ER- normal network topology changes drastically if different correlation cutoff values are used (**S. Fig. 2E, 2F**), we found that the cutoff 0.68 makes the ER- normal network most similar to the ER+ reference network (**S. Fig. 2G**). Using another measure for topological similarity, average clustering coefficient [33], the cutoff of 0.68 is supported again since normal ER- network with correlation cutoff 0.68 makes the closest average clustering coefficient to the reference network (0.4, **S. Fig. 2H**). Since normal and tumor ceRNA networks should share the cutoff within each subtype, we applied the subtype-specific cutoffs (0.68 for ER- and 0.6 for ER+) to build the tumor ceRNA networks.

### 3′UTR shortening is associated with the aggressive metastatic phenotypes of ER- tumors in ceRNA

In normal ER- ceRNA network based on the subtype-specific co-expression cutoff, 1,783 genes are in ceRNA relationship with 521 3′US genes (3′US ceRNA partners). Among 1,783 3′US ceRNA partners, 498 (27.9%) are found only in ER- (ER- 3′US ceRNA partners), whereas the other 1,285 (72.1%) are also in ER+ as 3′US ceRNA partners (common 3′US ceRNA partners, **Fig. 2A**). We found that 118 IPA canonical pathways significantly (P < 0.01) enriched for the ER- 3′US ceRNA partners (**S. Table 2**) are linked with several aspects of ER- specific tumor phenotypes (**Fig. 2B**). The first set of the pathways are “cancer” pathways. For example, the “Molecular Mechanisms of Cancer” pathway (P=10^−5.25^) includes a comprehensive set of genes, disruptions of which are known to promotes tumor growth. Specific to breast cancer, the enrichment of the “Breast Cancer Regulation by Stathmin1” (P=10^−3.92^) pathway is interesting, since overexpression of Stathmin1 correlates with loss of the ER [34] and with aggressive breast tumor phenotypes [35]. The second category of pathways underlies the aggressive metastasis of ER- tumors. For example, among eight pathways implicated in metastatic biology that were shown to play roles in breast tumor metastasis [36], we found that five of them are significantly enriched for ER- 3′US ceRNA partners with the exception of PI3K/AKT, the enriched p-value of which is just below the significance cutoff (P=10^−1.95^). Further, previous studies have associated breast tumor malignancy and poor survival with abnormal control of Ephrin A (reviewed in [37]), which is strongly enriched for ER- 3′US ceRNA partners (P-val=10^−5.05^). In normal samples without 3′-UTR shortening, 3′US ceRNA partners should closely regulate these pathways. However, in ER- tumors characterized by widespread 3′US events, most (81.7%) of the 3′US ceRNA partners lost the ceRNA relationship (**Fig. 2C)**, likely losing the normal control.

**Figure 2.**
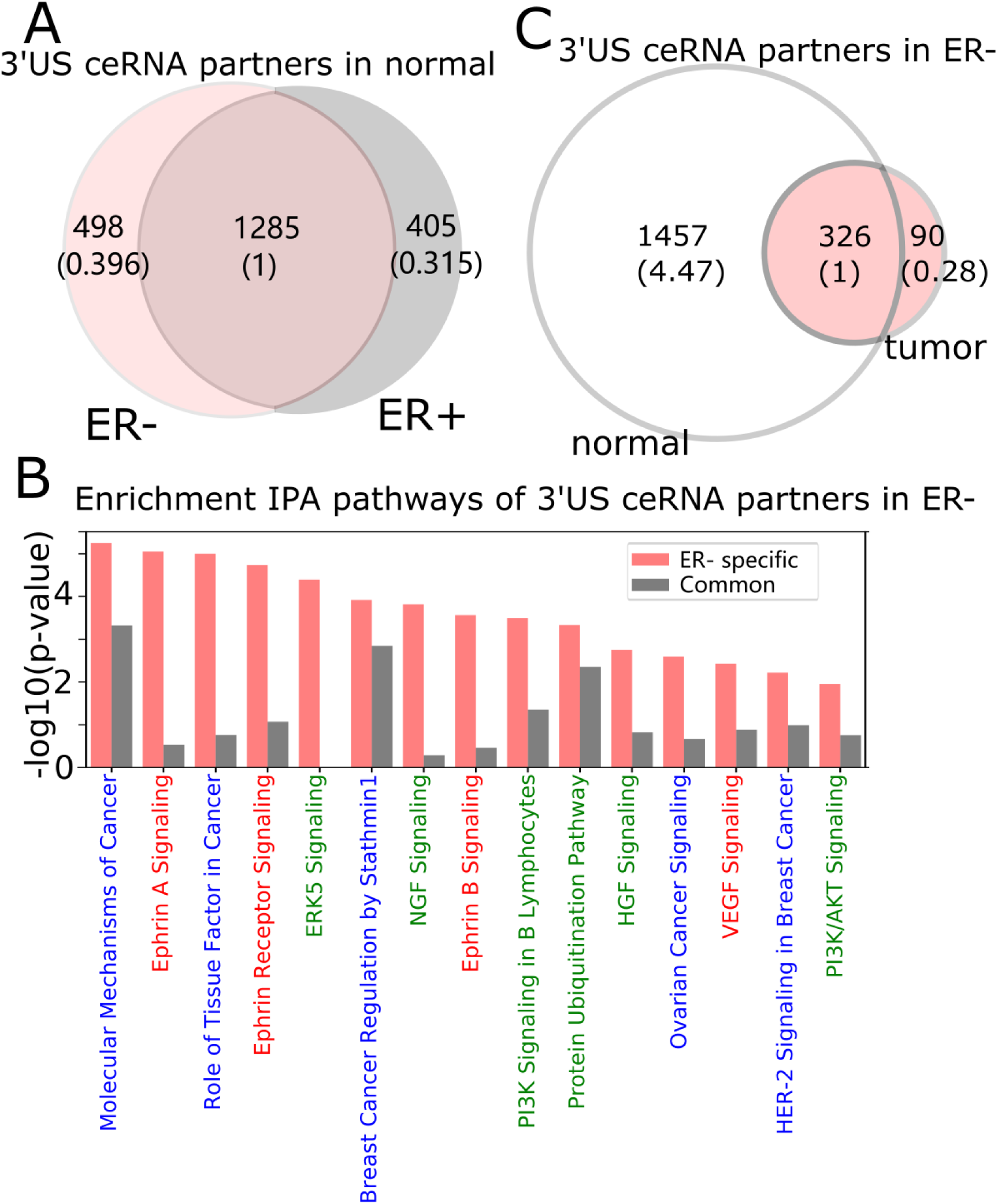
3′UTR shortening is associated to ER-’s aggressive phenotypes in ceRNA. (A) Intersection of 3′US ceRNA partners between ER- and ER+ normal ceRNA networks. (B) IPA canonical pathways significantly (P < 0.01) enriched for the ER- 3′US ceRNAs. Pathways are colorcoded by keyword, “Cancer” in blue, “Signaling” in red and those associated with aggressive phenotypes[25] in green. (C) Intersection of 3′US ceRNA partners in ER- between normal and tumor ceRNA networks.

### Housekeeping genes keep ER+ and ER- normal ceRNA networks to similar topology

Further, we categorized genes that have possible sponge effect (>5 miRNA binding sites in the 3′UTR) into housekeeping (HK), tumor-associated (tumor suppressors or oncogenes, TA), and transcription factor (TF). Based on 3,804 HK [26], 932 TA [28], and 1,020 TF genes [27] curated in public databases (see Methods), the ceRNA networks consist of 3-fold more HK genes than TA or TF genes (**Fig. 3A** for normal and **S. Fig. 3A** for tumor). Due to their active roles in cell maintenance [26], HK genes are expected to maintain constant expression levels under most physiological conditions [26]. Accordingly, the 958 HK ceRNA genes in ER- normal (**Fig. 3A**) express as highly as (**S. Fig. 3B)**, but with less significant variation (P=1.72e^−54^) across the normal samples (**Fig. 3B**), than 1,906 non-HK ceRNA genes in the network. With our observation that the HK genes contain more miRNA binding sites than the other genes (P=0.05, **Fig. 3C**), they should function as stable sponges for miRNAs [38]. Thus, with a significant number (P=8.77e^−771^) of overlap in the HK ceRNA genes (**Fig. 3D**) between ER- and ER+ normal samples, we hypothesize that they keep ER- and ER+ normal ceRNA networks in similar topology. To test this hypothesis, we first selected edges involving the HK ceRNA genes from the ER+ and ER- normal ceRNA networks to form subnetworks and compared the subnetworks using normalized Laplacian Matrix Eigenvalue Distribution. Further, we randomly subsampled the same number of edges not involving HK genes 200 times from the ER+ and ER- ceRNA networks and compare the networks in the same way. The HK ceRNA networks are significantly more similar between ER+ and ER- (P < 0.01) than 200 non-HK ceRNA networks, suggesting that HK genes make normal ceRNA crosstalk consistent between the subtypes through the miRNA sponge effect.

**Figure 3.**
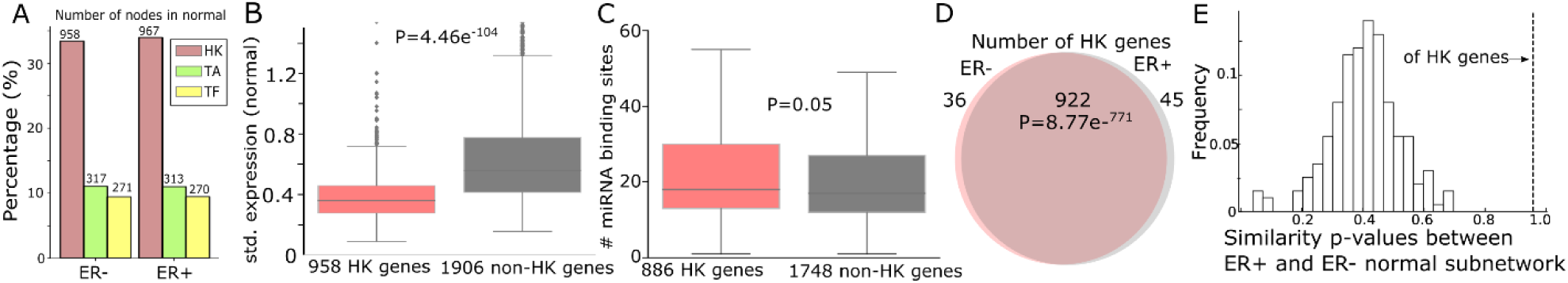
Housekeeping genes make consistent ceRNA networks between ER- and ER+ normal samples. (A) The number (and the percentage to the total number of nodes in the networks) of housekeeping (HK), tumor-associated (TA), or transcription factor (TF) genes in ER+ and ER- normal ceRNA networks. (B) Standard deviation of gene expressions of 958 HK genes and 1,906 non-HK genes in ER- normal ceRNA network. (C) Number of miRNA binding sites on the 3′UTR of 886 HK and 1,748 non-HK genes in the network (those that have miRNA binding site information). (D) Number of HK genes shared by ER- and ER+ normal ceRNA networks (with those in common). (E) Distribution of the similarity p-values between subnetworks sampled with 922 HK genes from ER+ and ER- normal networks (horizontal dotted line) and 200 subnetworks sampled with 1,990 non-HK genes to the same number of HK genes (bar). The higher the p-value is, the more similar the networks are [21].

### 3′US disrupts ceRNA crosstalk of housekeeping genes for ER- specific growth

We further examined the impact of 3′US on the role of HK genes. First, 3′US genes are highly connected to HK genes. Out of 958 HK genes, 727 HK genes (75.8%) are connected to 3′US genes, which is in the same scale as the other classes of genes that are known to be regulated by 3′US genes [10], [12] (61.8% from 317 TA genes and 90.2% from 271 TF genes, **Fig. 4A**). Also, these HK genes are more highly connected in the network (**Fig. 4B**) compared to 231 HK genes that are not connected to 3′US genes. Previously, we showed that 3′US represses the ceRNA partners in tumor [12]. Consistently, these HK genes, ceRNA partners of 3′US genes, are more repressed in tumor than 231 HK genes not connected to 3′US genes (**Fig. 4C**). To understand the impact of the repression on the ceRNA network, we compared the number of the ceRNA partners of these HK genes between normal and tumor. Previously, we showed that 3′US genes will break their relationship with the ceRNA partners [12]. Since the ceRNA relationship changes, either loss or gain, could propagate to neighboring ceRNA relationships [11], the repression of HK genes should break the ceRNA relationship not only with 3′US genes but also with other ceRNA partners. Consistent to the expectation, 727 3′US HK ceRNA partners lost higher ratios of the ceRNA partners in tumor (**Fig. 4D**).

**Figure 4.**
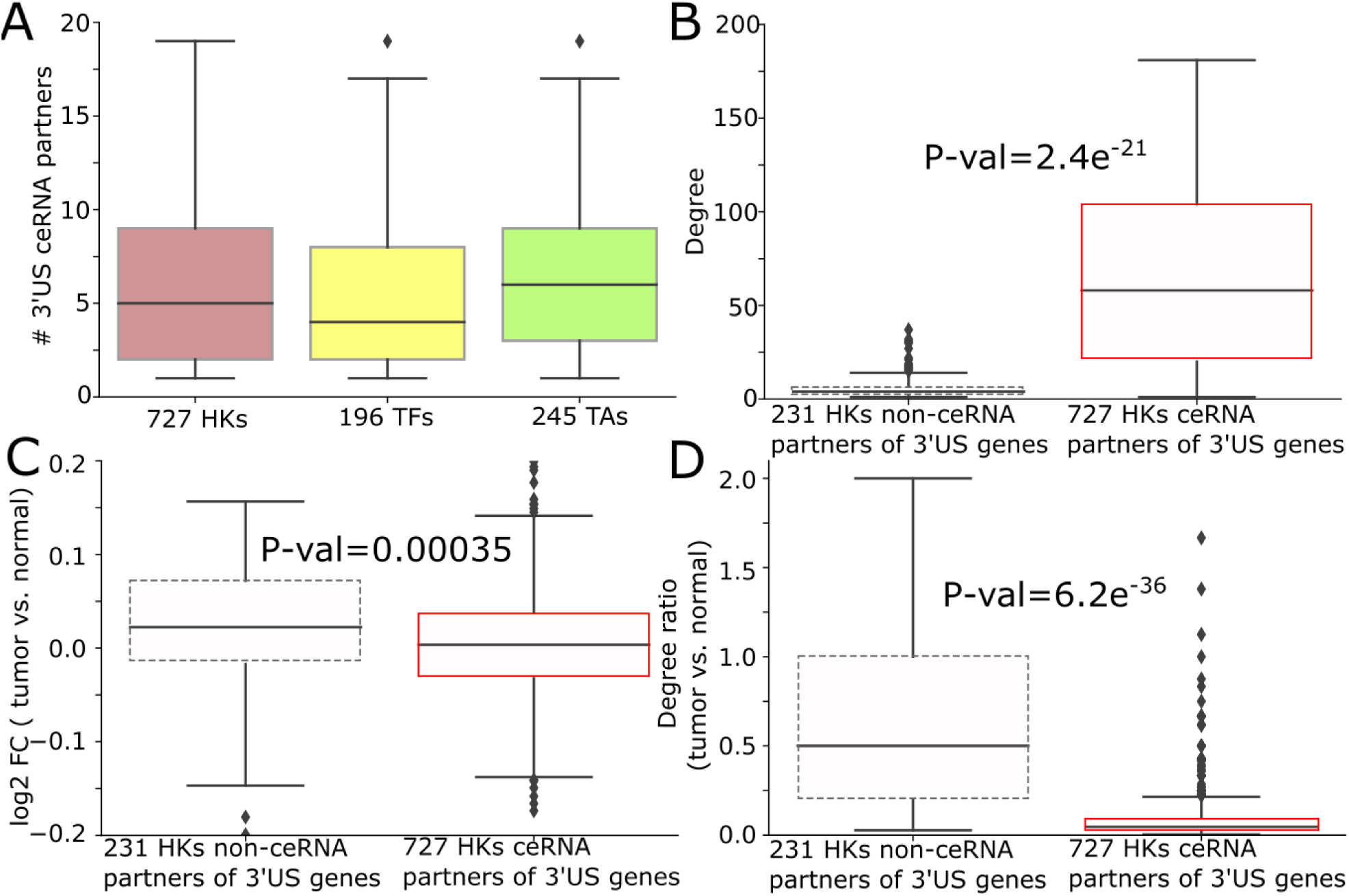
3′US disrupts ceRNA relationship of HK genes. (A) # of 3′US genes connected to housekeeping (HK), transcription factor (TF), and tumor-associated (TA) genes (B) Degree (# neighbors in ERN normal ceRNA network), (C) log2 fold change (tumor vs. normal), (D) degree ratio (tumor vs. normal) of 727 and 231 HK genes that are ceRNA partners of 3′US genes or not, respectively. Since degree ratio in (D) represents the ratio of the number of neighbors retained in tumor, low degree values of 727 3′US HK ceRNA partners represents their higher loss of ceRNA neighbors in tumor.

The loss of HK ceRNA partners naturally reduces the high overlap of HK genes between ER+ and ER- (**Fig. 5A**), resulting into 505 and 144 HK genes that are ceRNA partners of 3′US genes unique in ER- and ER+ tumor (ER- and ER+ HK ceRNA partners), respectively (**Fig. 5B**). While it is known that cell growth and cell cycle regulations are different in the subtypes [39]– [41], we found that 505 ER- HK ceRNA partners are enriched for cell growth- and cell cycle-related IPA pathways. First, they are enriched for pathways associated to growth factor (with keyword “GF”) (**Fig. 5C, S. Table 4**). Especially, EGF (P-val=10^−2.99^) activates cell cycle progression in ER- tumors [42], and expression of VEGF (P-val=10^−2.42^) is associated to ER- tumors [43]. Also, both EGF and VEGF are suspected to proliferate ER- tumors when estrogen cannot sustain them [43]. Second, cell cycle pathways are enriched for ER+ specific HK ceRNA partners, suggesting that ER-regulated cell cycle [31], [32] differentiates ER+ and ER- cancer partially at the ceRNA level. Especially, since regulation of cell cycle, G1- and S-phase and their transition ratio, is crucial for ER+ tumor’s proliferation (reviewed in [44]), it is interesting that cell cycle regulation pathways for various phases (G1/S or G2/M) of various mediators (Estrogen or Cyclins) are enriched with 144 ER+ HK ceRNA partners. Third, considering that the enrichment analysis was for the disjoint sets of genes (505 unique to ER- and 144 unique to ER+), it is interesting that these unique HK ceRNA partners are commonly significantly enriched for some “cancer” pathways e.g. “Molecular Mechanisms of Cancer”, showing that the HK ceRNAs are involved in cancer mechanisms equally significantly but in a subtype-specific fashion.

**Figure 5.**
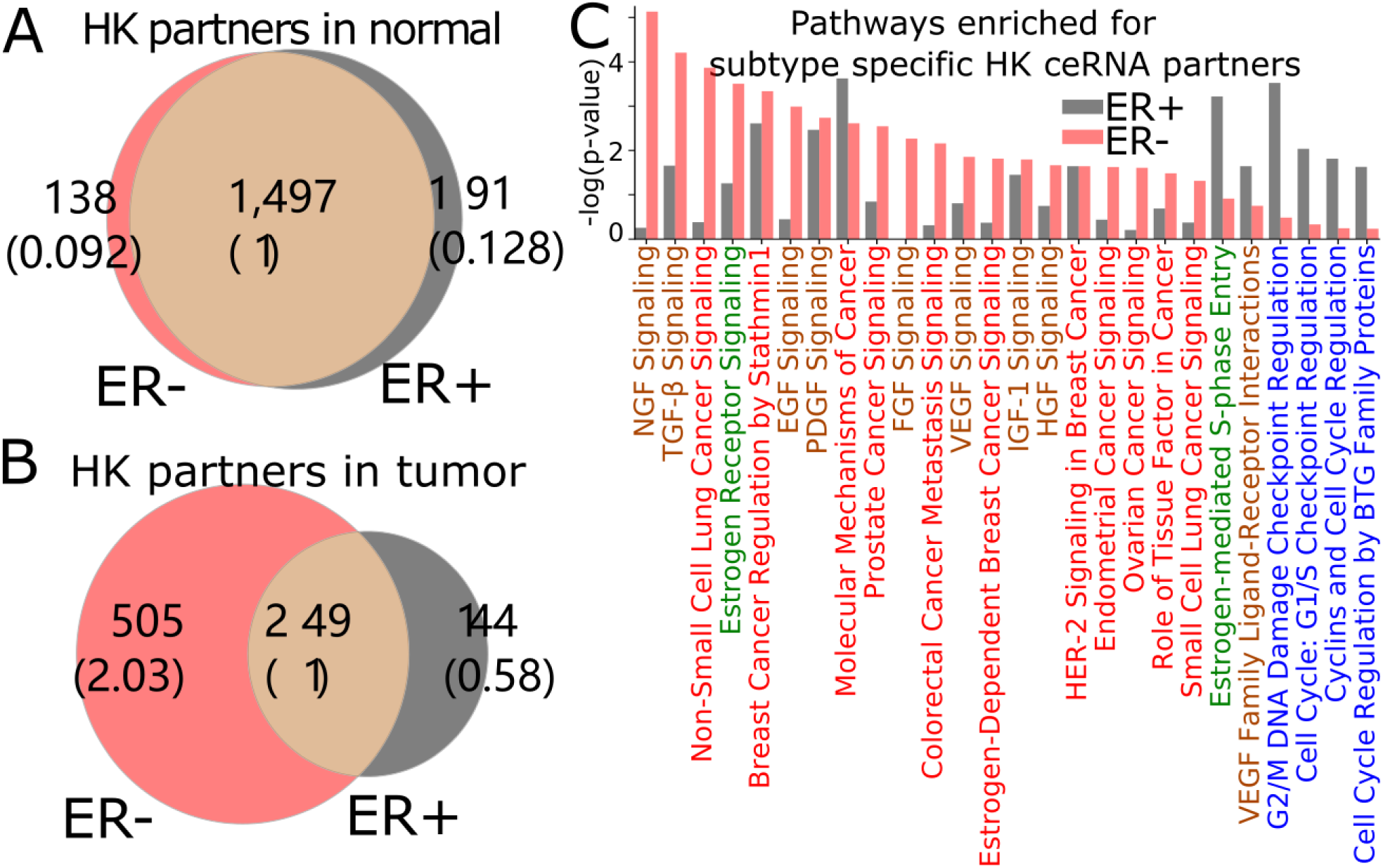
3′US disrupts ceRNA relationship of HK genes for ER- specific growth. (A) Number of HK ceRNA partners unique and common to ER- and ER+ normal (left) and tumor (right) ceRNA networks. The numbers in parentheses are normalized to the number of genes shared between tumor and normal. (B) Degree in ER- tumor ceRNA network of 727 and 231 HK genes that are ceRNA partners of 3′US genes or not, respectively (C) IPA canonical pathways significantly (P < 0.01) enriched for ER+ and ER- specific HK ceRNA partners. Pathways are color-coded by keyword, “Cancer” in red, “GF” in brown, “Estrogen” in green, and “Cell Cycle” in blue.

### 3′US represses housekeeping genes to promote tumor growth

To gain insights into the cause-and-effect relationship from 3′US-mediated HK gene repression to tumorigenesis, we revisited a previous study [12], [45], in which 3′US-ceRNA effect promotes tumorigenesis in NUDT21 knockdown (KD) in HeLa cells and glioblastoma (data available in GSE42420 [14] and GSE78198 [31]). First, we chose 11,431 genes that are expressed in the experiment data (avg. FPKM > 1). Among them, we further chose 4,430 genes that would work as miRNA sponges (>5 miRNA binding sites). To identify ceRNA relationship with the genes, we will solely use significance of miRNA binding site overlap (FDR < 0.05), since the other criteria for the ceRNA identification, co-expression, cannot be effectively estimated from two replicates of NUDT21 KD experiments. In this way, we identified 860 3′US genes and 2,449 of their ceRNA partners. Among these 3′US ceRNA partners, a significant portion of them (705, 28.8%) are HK genes, while 184 are TA and 163 are TF genes. Especially, it is interesting to note that HK genes in the network are only either 3’US genes (n=298) or 3’US ceRNA partners (n=705). On the other hands, almost half of the TA and TF genes in the network are not connected with 3’US genes (149 of 333 (44.7%) and 147 of 310 (47.4%) for TA and TF, respectively), showing that HK genes can be a major target of 3’US ceRNA effect. Based on our previous finding that 3′US represses the ceRNA partners in tumor [12], we further checked the repression of HK genes in NUDT21 KD. 705 HK genes that are 3′US ceRNA partners are more repressed than TA and TF genes or than 298 HK 3’US genes in the network (**Fig. 6A**, P-value=0.01 and 0.05, 0.002, respectively). These results confirm that HK genes are repressed in the tumorigenic process 3′US-ceRNA effect promotes [12].

**Figure 6.**
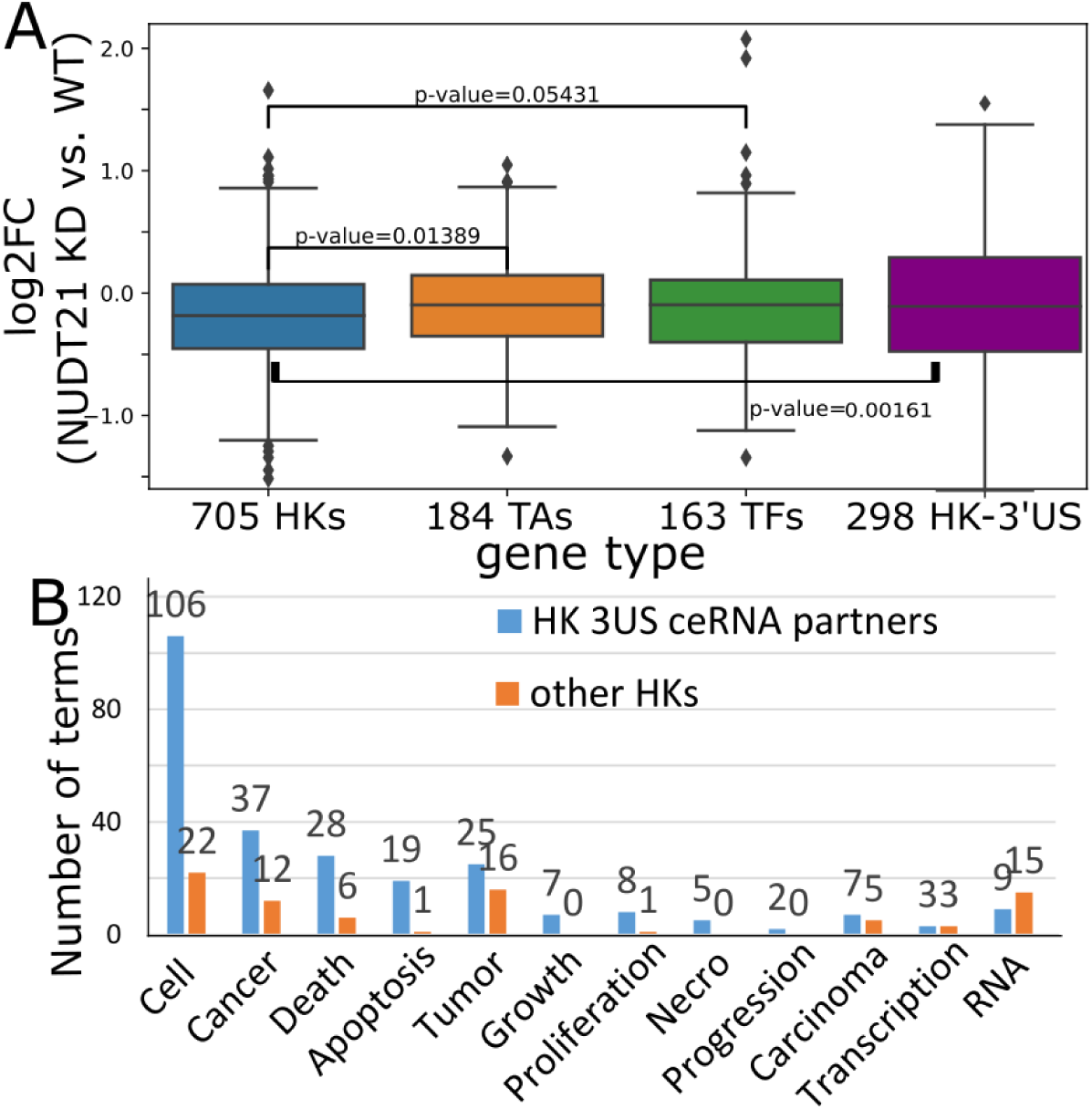
(A). log2FC (NUDT21 KD vs. WT) of 705 HK, 184 TA, and 163 TF genes that are (potential) ceRNA partners of 3’US genes. log2FC of 298 HK genes that are 3’US genes on the rightmost box. (B). Number of terms with the keyword indicated on the x-axis. Numbers on bar represent the actual number of terms.

To assess the impact of this repression on tumor growth, we further conducted IPA analysis on 705 HK 3’US ceRNA partners in comparison to the other 2,410 HK genes not in the network. First, although there are much less HK 3’US ceRNA partners than the other HK genes, they are enriched for more IPA Diseases & Functions terms (**S. Table 4**). While the IPA analysis gives N/A for the terms that are so lowly enriched that cannot be estimated, HK 3’US ceRNA partners have 581 terms with N/A value and HK genes not in the network have 693 terms with N/A value. Further, we replaced the N/A values with the minimum value and compare the p-values in HK 3’US ceRNA partners vs. the other HK genes. This comparison shows that more terms are significantly (P-value < 0.01) enriched for HK 3’US ceRNA partners (254 terms with better p-values for HK 3’US ceRNA partners and 141 for the other HK genes). This trend is more pronounced for the terms that are important for cancer. For example, IPA terms with keywords “Cell”, “Cancer (or Tumor)”, “Apoptosis (, Death, or Necro)”, and “Growth (, Proliferation, or Progression)” are significantly (P-value < 2.2e^−16^) more enriched in the HK 3’US ceRNA partners, while certain terms for general biological processes such as “RNA” are enriched in the other HK genes (**Fig. 6B**). Altogether, the results show that, when 3′US-ceRNA effect promotes tumor growth in NUDT21 KD experiment [12], it does so partly by repressing HK genes.

## DISCUSSION

To investigate the role of 3′US-ceRNA effect [12] for estrogen receptor negative (ER-) vs. ER+ breast tumors, we built the ceRNA networks that are comparable to each other subtype by addressing the bias due to the different number of samples (72 for ER+ and 20 for ER- in TCGA). A fair comparison of the networks suggests that 3′US disrupts the ceRNA network for ER- tumors’ aggressive phenotypes. Further, we revealed a role of 3′US-ceRNA effect on housekeeping (HK) genes. Although HK genes highly and stably express in diverse biological contexts [25], our understanding of their roles is limited, especially with regards to ceRNA crosstalk. For the first time, our results identified a novel role of HK genes in maintaining the ceRNA network on physiological conditions with normal samples adjacent to tumor samples as an example.

Further analyses show that 3′US disrupts ceRNA crosstalk of HK genes in a subtype-specific fashion. First, we showed that a subset of HK genes is *trans* target of 3′US-ceRNA effect (sponge HK genes) enriched in important pathways in association to ER-’s aggressive phenotype. Since they are much less than the other HK genes in number (e.g. 705 3′US ceRNA HK genes vs. 2,401 HK genes in the NUDT21 KD experiment), our definition may shed novel insights into identifying another set of biomarkers indicating tumor progression.

In network analysis, a network of interest is often compared to a reference network. However, if the networks are built from different numbers of samples, the comparison will be biased due to the sample size difference (**S. Fig. 2**). With the assumption that normal samples should have similar molecular dynamics, we found the subtype-specific cutoff values for normal ceRNA networks. Then, we assumed that normal and tumor ceRNA network should share the cutoff within each subtype to construct ER+ and ER- tumor ceRNA networks (two-step pairwise normalization method). As the resulting ceRNA networks facilitate novel discoveries on the subtype-specific 3′US-ceRNA effect, we expect that the two-step pairwise normalization method can further help normalize biological networks built with the different number of samples if the matched normal samples are available.

---

**S. Figure 1.**
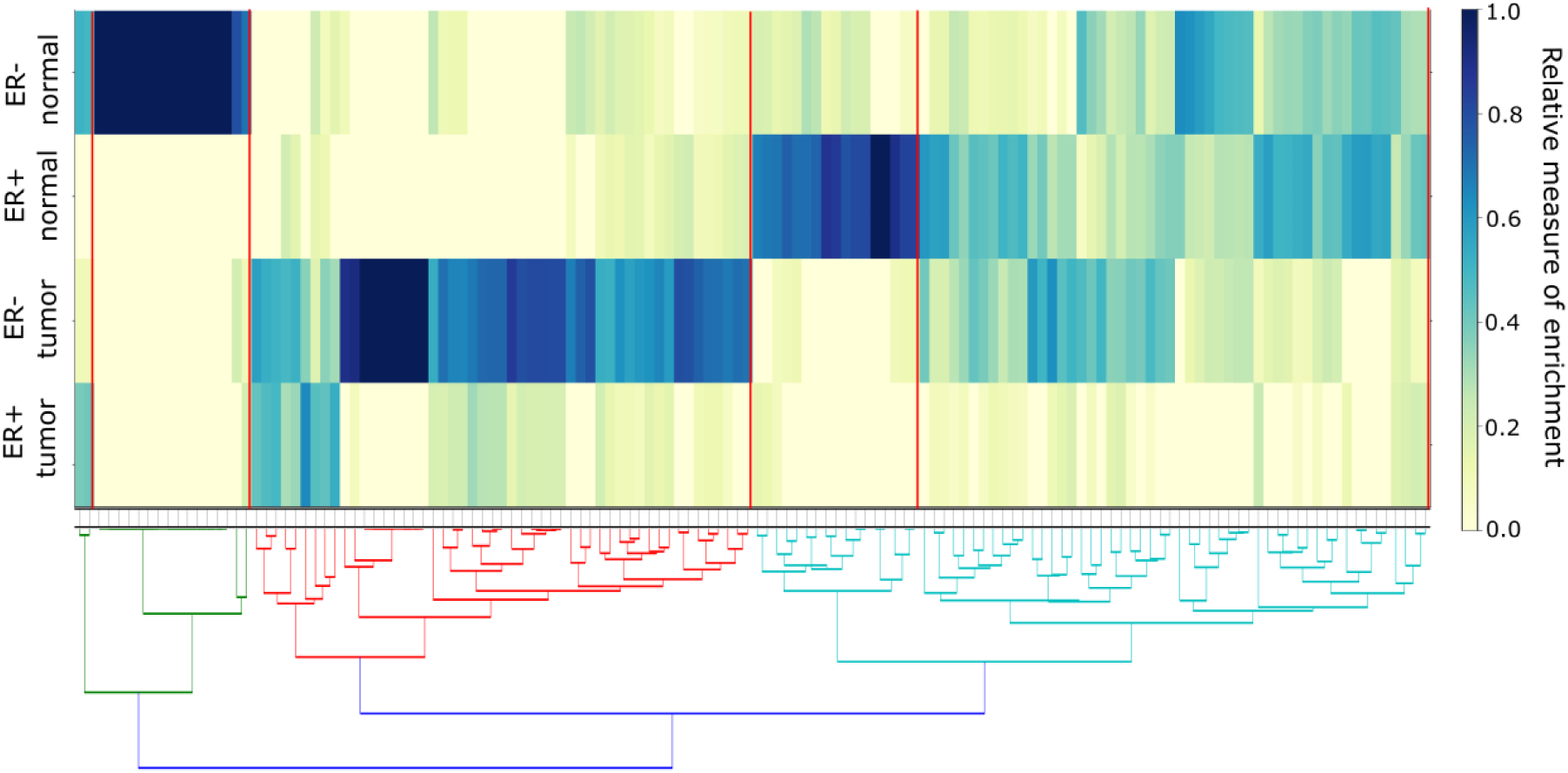
IPA pathways enriched for 3′UL and 3′US genes in ER- and ER+. Colors represent enrichment of each pathway (column) for each class of genes (The higher the enrichment is, the higher the associated term is enriched). The red lines cut the pathways into 5 clusters, where each cluster is enriched in a set of genes.

**S. Figure 2.**
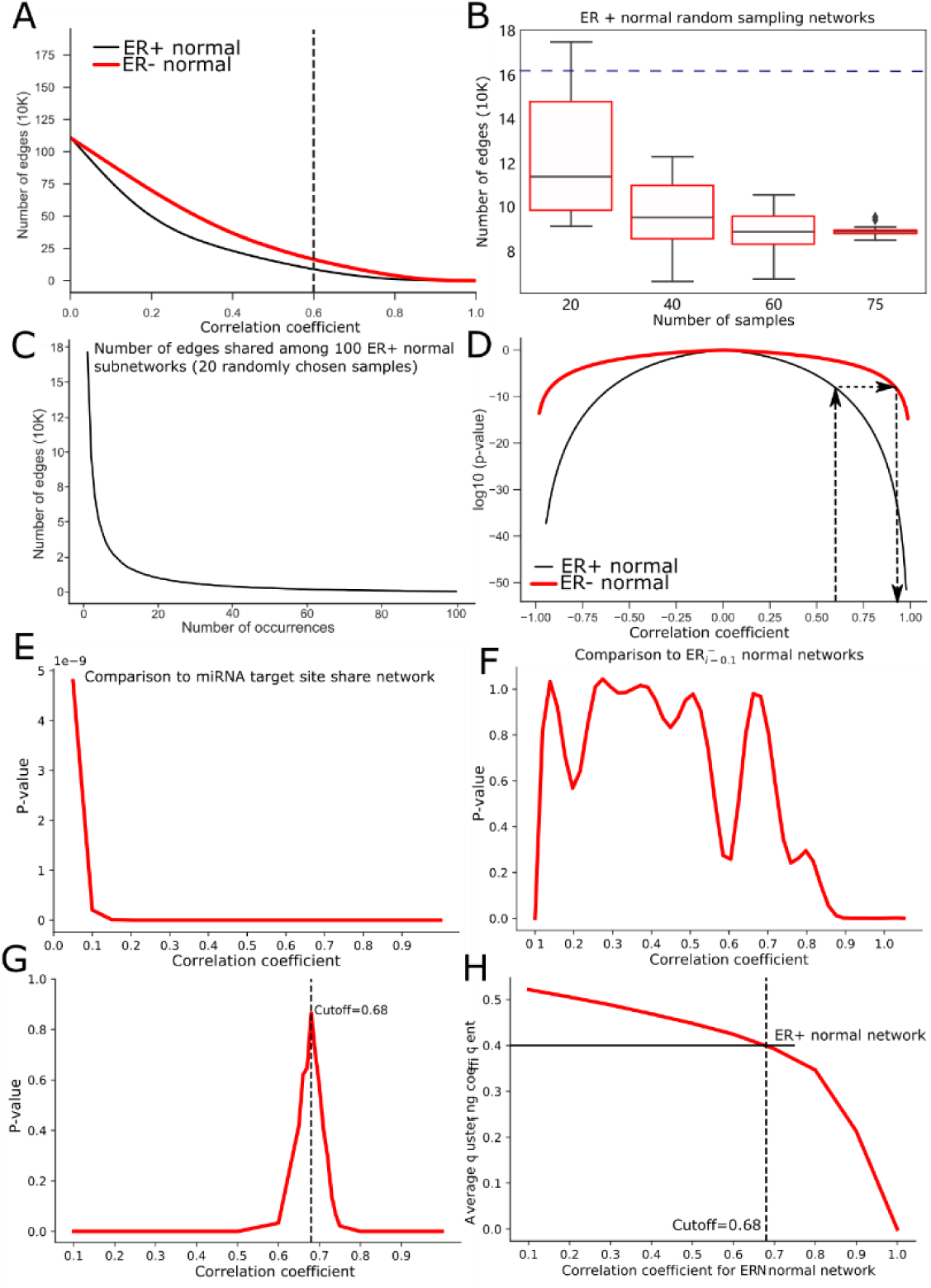
Two-step Pairwise Normalization of ER+ and ER- ceRNA network. (A) Number of edges in the ceRNA networks by the correlation coefficient cutoff (black and red line for ER+ and ER- networks, respectively). (B) Number of edges of networks from a subset of ER+ normal samples sampled in different size. Blue dotted line represents the number of edges of ER- normal network whose sample size is 20 (160,687) (C) The number of edges shared among 100 ER+ subnetworks from normal samples, where each of them was built by using 20 randomly chosen samples. (D) Statistical significance (p-value) achievable by using different correlation coefficient cutoff values for ER+ (black) and ER- (red) samples. Statistical significance for a correlation coefficient cutoff value is described in Methods. To achieve the same statistical significance of the traditional cutoff value (0.6) of ER+ in ER-, the cutoff value would inflate to 0.91, resulting in drastically a deflated number of edges. (E) Comparison of the miRNA target site share network to ER- normal networks with different correlation cutoff values (see Methods). As illustrated, ER- normal network topology changes drastically by different cutoff values in reference to the miRNA target site share network. (F) Comparison of ER- normal networks with previous correlation cutoff values in the stepwise increase (see Methods). (G). Topological similarity (y-axis) between ER+ and ER- normal ceRNA networks by the cutoff value for ER- (x-axis). The bigger the p-value is, the more similar the two networks are (see Methods)[21]. The ER- normal network with the cutoff of 0.68 looks most similar to the ER+ normal reference network. (H) Comparing ER- normal ceRNA networks of different correlation cutoff values with the ER+ normal reference network in the average clustering coefficient. The average clustering coefficient for the ER+ normal reference network is 0.40 (indicated by the horizontal black line), which is quite close to the average clustering coefficient for the ER- normal network with a cutoff of 0.68 (indicated by the vertical dashed line).

**S. Figure 3.**
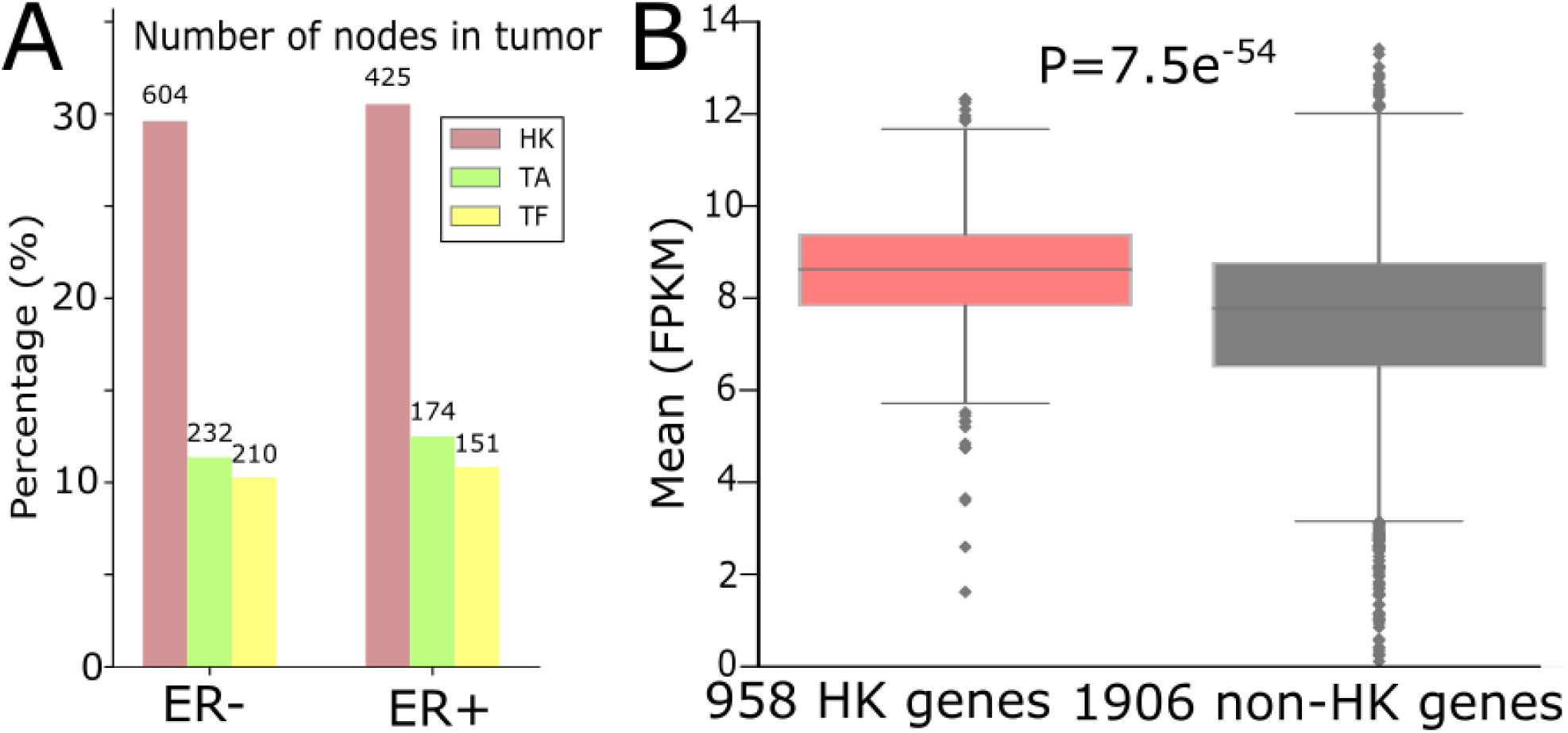
(A) Number (and percentage to the total number of nodes in tumor networks) of HK genes and other important classes of genes in ER+ and ER- normal ceRNA networks. (B) Average gene expression values of 958 HK genes and 1,906 non-HK genes in ER+ and ER- normal networks.

## Acknowledgements

We thank Susanne M. Gollin, Ph.D., Professor of the Department of Human Genetics in the University of Pittsburgh for their valuable help in editing this manuscript. Also, this research was supported in part by the University of Pittsburgh Center for Research Computing through the resources provided. This manuscript has been released as a Pre-Print at [46].

## Competing interests

The authors declares no competing financial interests.

## Funding

This work was supported partly by CTSI Biomedical Modeling Awards and the Joan Gollin Gaines Cancer Research Fund at the University of Pittsburgh to H.J.P..

## Author Contributions

H.J.P conceived the project and designed the experiments. F.Z. performed the data analysis. S.K. conducted statistical tests. H.J.P. wrote the manuscript with input from B.D., S.K..

## Notes

#### Summary of Updates

It was updated to highlight the importance of our novel findings on sponge housekeeping genes.

## REFERENCES

[1] C. Mayr and D. P. Bartel, “Widespread shortening of 3’UTRs by alternative cleavage and polyadenylation activates oncogenes in cancer cells.,” Cell, vol. 138, no. 4, pp. 673–84, Aug. 2009.

[2] M. Chen et al., “3 ′ UTR lengthening as a novel mechanism in regulating cellular senescence,” pp. 285–294, 2018.

[3] G. P. Dimri et al., “A biomarker that identifies senescent human cells in culture and in aging skin in vivo.,” Proc. Natl. Acad. Sci. U. S. A., vol. 92, no. 20, pp. 9363–7, 1995.

[4] R. A. Busuttil, M. Rubio, M. E. T. Dollé, J. Campisi, and J. Vijg, “Oxygen accelerates the accumulation of mutations during the senescence and immortalization of murine cells in culture.,” Aging Cell, vol. 2, no. 6, pp. 287–294, 2003.

[5] C. López-Otín, M. A. Blasco, L. Partridge, M. Serrano, and G. Kroemer, “The hallmarks of aging,” Cell, vol. 153, no. 6, 2013.

[6] D. Muñoz-Espín and M. Serrano, “Cellular senescence: From physiology to pathology,” Nat. Rev. Mol. Cell Biol., vol. 15, no. 7, pp. 482–496, 2014.

[7] Z. Xia et al., “Dynamic Analyses of Alternative Polyadenylation from RNA-Seq Reveal Landscape of 3’ UTR Usage Across 7 Tumor Types,” Nat. Commun., pp. 1–38, 2014.

[8] Y. Xiang et al., “Comprehensive Characterization of Alternative Polyadenylation in Human Cancer,” vol. 110, no. November 2017, pp. 1–11, 2018.

[9] L. Salmena, L. Poliseno, Y. Tay, L. Kats, and P. P. Pandolfi, “A ceRNA hypothesis: the Rosetta Stone of a hidden RNA language?,” Cell, vol. 146, no. 3, pp. 353–8, Aug. 2011.

[10] P. Sumazin et al., “An extensive microRNA-mediated network of RNA-RNA interactions regulates established oncogenic pathways in glioblastoma.,” Cell, vol. 147, no. 2, pp. 370–81, Oct. 2011.

[11] H. J. Park, S. Kim, and W. Li, “Model-based analysis of competing-endogenous pathways (MACPath) in human cancers,” PLoS Comput. Biol., vol. 22, no. 14, 2018.

[12] H. J. Park et al., “3′ UTR shortening represses tumor-suppressor genes in trans by disrupting ceRNA crosstalk,” Nat. Genet., vol. 50, pp. 783–789, 2018.

[13] S. Kim, Y. Bai, Z. Fan, B. Diergaarde, G. C. Tseng, and H. J. Park, “Alternative Polyadenylation Regulates Patient-specific Tumor Growth by Individualizing the MicroRNA Target Site Landscape,” bioRxiv, p. 601518, Jan. 2019.

[14] M. E. H. Hammond et al., “American Society of Clinical Oncology/College of American Pathologists guideline recommendations for immunohistochemical testing of estrogen and progesterone receptors in breast cancer (unabridged version).,” Arch. Pathol. Lab. Med., vol. 134, no. 7, pp. e48–72, Jul. 2010.

[15] S. Tsutsui, S. Ohno, S. Murakami, Y. Hachitanda, and S. Oda, “Prognostic value of epidermal growth factor receptor (EGFR) and its relationship to the estrogen receptor status in 1029 patients with breast cancer,” Breast Cancer Res. Treat., vol. 71, no. 1, pp. 67–75, 2002.

[16] M. Sheikh, G. M, P. P, F. JA, and R. H, “Why are estrogen-receptor-negative breast cancers more aggressive t,” Invasion Metastasis, vol. 14, no. 1–6, pp. 329–36, 1994.

[17] A. Pergamenschikov et al., “Molecular portraits of human breast tumours,” Nature, vol. 406, no. 6797, pp. 747–752, 2002.

[18] M. Goldman et al., “The UCSC Cancer Genomics Browser: update 2013.,” Nucleic Acids Res., vol. 41, no. Database issue, pp. D949–54, Jan. 2013.

[19] B. P. Lewis, C. B. Burge, and D. P. Bartel, “Conserved seed pairing, often flanked by adenosines, indicates that thousands of human genes are microRNA targets.,” Cell, vol. 120, no. 1, pp. 15–20, Jan. 2005.

[20] G. L. Papadopoulos, M. Reczko, V. a Simossis, P. Sethupathy, and A. G. Hatzigeorgiou, “The database of experimentally supported targets: a functional update of TarBase.,” Nucleic Acids Res., vol. 37, no. Database issue, pp. D155–8, Jan. 2009.

[21] F. Xiao, Z. Zuo, G. Cai, S. Kang, X. Gao, and T. Li, “miRecords: An integrated resource for microRNA-target interactions,” Nucleic Acids Res., vol. 37, no. November 2008, pp. 105–110, 2009.

[22] S.-D. Hsu et al., “miRTarBase update 2014: an information resource for experimentally validated miRNA-target interactions.,” Nucleic Acids Res., vol. 42, no. Database issue, pp. D78–85, Jan. 2014.

[23] H. Dvinge et al., “The shaping and functional consequences of the microRNA landscape in breast cancer.,” Nature, vol. 497, no. 7449, pp. 378–82, May 2013.

[24] M. P. Hamilton et al., “Identification of a pan-cancer oncogenic microRNA superfamily anchored by a central core seed motif.,” Nat. Commun., vol. 4, p. 2730, Jan. 2013.

[25] E. Eisenberg and E. Levanon, “Human housekeeping genes are compact,” TRENDS Genet., vol. 19, no. 7, pp. 362–365, 2003.

[26] E. Eisenberg and E. Y. Levanon, “Human housekeeping genes, revisited.,” Trends Genet., vol. 29, no. 10, pp. 569–74, Oct. 2013.

[27] K. Chawla, S. Tripathi, L. Thommesen, A. Lægreid, and M. Kuiper, “TFcheckpoint: A curated compendium of specific DNA-binding RNA polymerase II transcription factors,” Bioinformatics, vol. 29, no. 19, pp. 2519–2520, 2013.

[28] T. Davoli et al., “Cumulative Haploinsufficiency and Triplosensitivity Drive Aneuploidy Patterns and Shape the Cancer Genome.,” Cell, vol. 155, no. 4, pp. 948–962, Oct. 2013.

[29] U. Ala et al., “Integrated transcriptional and competitive endogenous RNA networks are cross-regulated in permissive molecular environments.,” Proc. Natl. Acad. Sci. U. S. A., vol. 110, no. 18, pp. 7154–9, Apr. 2013.

[30] R. Gera et al., “Identifying network structure similarity using spectral graph theory,” Appl. Netw. Sci., vol. 3, no. 1, p. 2, 2018.

[31] S. Javanmoghadam, Z. Weihua, K. K. Hunt, and K. Keyomarsi, “Estrogen receptor alpha is cell cycle-regulated and regulates the cell cycle in a ligand-dependent fashion,” Cell Cycle, vol. 15, no. 12, pp. 1579–1590, 2016.

[32] S. Paruthiyil, H. Parmar, V. Kerekatte, G. R. Cunha, G. L. Firestone, and D. C. Leitman, “Estrogen Receptor □ Inhibits Human Breast Cancer Cell Proliferation and Tumor Formation by Causing a G 2 Cell Cycle Arrest,” pp. 423–428, 2004.

[33] C. C. Friedel and R. Zimmer, “Inferring topology from clustering coefficients in protein-protein interaction networks,” BMC Bioinformatics, vol. 15, pp. 1–15, 2006.

[34] P. Curmi et al., “Overexpression of stathmin in breast carcinomas points out to highly proliferative tumours,” Br. J. Cancer, vol. 82, no. 1, pp. 142–150, 2000.

[35] S. Obayashi et al., “Stathmin1 expression is associated with aggressive phenotypes and cancer stem cell marker expression in breast cancer patients,” Int. J. Oncol., vol. 51, no. 3, pp. 781–790, 2017.

[36] K. Krishnan et al., “miR-139-5p is a regulator of metastatic pathways in breast cancer.,” RNA, vol. 19, no. 12, pp. 1767–80, Dec. 2013.

[37] D. Vaught, D. M. Brantley-Sieders, and J. Chen, “Eph receptors in breast cancer: Roles in tumor promotion and tumor suppression,” Breast Cancer Research. 2008.

[38] Y. Tay, J. Rinn, and P. P. Pandolfi, “The multilayered complexity of ceRNA crosstalk and competition,” Nature, vol. 505, no. 7483, pp. 344–352, Jan. 2014.

[39] G. Hong et al., “Genes Dysregulated to Different Extent or Oppositely in Estrogen Receptor-Positive and Estrogen Receptor-Negative Breast Cancers,” PLoS One, vol. 8, no. 7, p. e70017, 2013.

[40] M. C. Alles et al., “Meta-Analysis and Gene Set Enrichment Relative to ER Status Reveal Elevated Activity of MYC and E2F in the ‘Basal’ Breast Cancer Subgroup,” PLoS One, vol. 4, no. 3, p. e4710, 2009.

[41] M. C. Abba et al., “Gene expression signature of estrogen receptor α status in breast cancer,” BMC Genomics, vol. 6, pp. 1–13, 2005.

[42] D. K. Biswas, a. P. Cruz, E. Gansberger, and a. B. Pardee, “Epidermal growth factor-induced nuclear factor kappa B activation: A major pathway of cell-cycle progression in estrogen-receptor negative breast cancer cells,” Proc. Natl. Acad. Sci., vol. 97, no. 15, pp. 8542–8547, Jul. 2000.

[43] D. Fuckar et al., “VEGF expression is associated with negative estrogen receptor status in patients with breast cancer,” Int. J. Surg. Pathol., vol. 14, no. 1, pp. 49–55, 2006.

[44] D. C. Henley, J. S. Foster, J. Wimalasena, P. Seth, and A. Bukovsky, “Multifaceted Regulation of Cell Cycle Progression by Estrogen: Regulation of Cdk Inhibitors and Cdc25A Independent of Cyclin D1-Cdk4 Function,” Mol. Cell. Biol., vol. 21, no. 3, pp. 794–810, 2002.

[45] C. P. Masamha et al., “CFIm25 links alternative polyadenylation to glioblastoma tumour suppression.,” Nature, vol. 510, no. 7505, pp. 412–416, May 2014.

[46] F. Zhenjiang, S. Kim, B. Diergaarde, and H. J. Park, “3′-UTR shortening contributes to subtype-specific cancer growth by breaking stable ceRNA crosstalk of housekeeping genes,” bioRxiv, p. 601526, Jan. 2019.

